# A patient-derived ovarian cancer organoid platform to study susceptibility to natural killer cells

**DOI:** 10.1101/2025.03.06.641285

**Authors:** Marisa Mercadante, Armin Scheben, Jacob Estrada, Jan Savas-Carstens, William Sullivan, Nicholas Housel, Tatiana Volpari, Jax Hebner, Maria Sapar, Tom Rusielewicz, Frederick J. Monsma, Stefan Semrau, Yinan Wang, Laura A. Martin

## Abstract

Intratumoral heterogeneity drives therapy resistance and relapses in advanced stage cancers, such as ovarian cancer. Here, we present a live cell imaging assay using patient-derived ovarian cancer organoids for real time capture and quantification of natural killer cell-mediated apoptotic events in >500 organoids simultaneously. Our assay revealed significant inter- and intratumor response heterogeneity and identified a rare resistant organoid population, opening avenues to test immunomodulatory strategies that overcome resistance.

## Introduction

Inter- and intratumor heterogeneity, which characterizes metastatic disease, drive resistance to therapies and most cancer-related deaths^1–5^. This heterogeneity also hampers recently developed immunotherapies that are revolutionizing cancer treatment for certain patients, including those with advanced stage disease. But despite the unprecedented efficacy of immunotherapies for hematological cancers and some patients with intractable tumors, they remain ineffective for most patients across many solid cancer types. A major roadblock for developing effective immunotherapies for heterogeneous cancers is characterizing patient-specific tumor-immune cell interactions that are driving the diverse array of responses, including resistance.

Among solid cancers, ovarian cancer represents one of the most heterogenous cancer types. Most patients are diagnosed with metastatic disease, where tumors arise from divergent clonal trajectories and present heterogeneous immune cellularity and immune escape mechanisms^6–10^. Subclonal resistance mechanisms underlie advanced stage disease with metastatic tumors exhibiting reduced immune cell infiltration and immunosuppressive microenvironments that are subtype-, patient- and even tumor site–specific^8–14^. T cell-based therapies and immune checkpoint inhibitors have so far been ineffective in treating ovarian cancer^15,16^. In contrast, positive results have been reported in clinical trials testing the efficacy of natural killer (NK) cells, including for heavily treated patients with refractory ovarian cancer^17–23^. Allogeneic NK cells are emerging as a promising alternative as they present a better safety profile than CAR-T cells and exhibit cytotoxic activity that is independent of specific surface target antigens, which might have advantages in heterogeneous tumors^24–26^. While primary NK cells have typically been used as a source, the recent availability of induced pluripotent stem cells (iPSCs)-derived NK cells (iNKs) provides advantages by ensuring uniform quality, being amenable to genetic engineering and allowing production at scale. Thus, clinical grade iNKs are in development to exploit them as a potential “off-the-shelf” cell therapy product. However, there is a need to evaluate their efficacy preclinically to accelerate clinical development. Most importantly, it is critical to identify tumors that present resistance and screen for strategies to overcome it.

Co-culture systems using cancer cells grown as monolayer or spheroids have been used to evaluate the potential therapeutic activity of NK cells. While these systems provide useful technical and biological insights, they lack the relevance of primary patient material and results are therefore poorly translatable to heterogenous advanced tumor types and patient populations^27–29^. In contrast, patient-derived tumor organoids (PDOs) are emerging as an invaluable system for studying immune-cancer cell interactions across several solid tumor types and optimizing combinatorial immunotherapy strategies^27,30–38^. PDOs maintain the complex 3D tissue architecture of primary tumors and recapitulate individual tumor phenotypes, including inter- and intratumor drug response heterogeneity^39–43^ and patient responses to chemotherapies^41,44–50^. In a recent study, PDOs in co-culture with tumor-infiltrating lymphocytes were established as a scalable immunotherapeutic modeling system suitable for combinatorial therapy testing^37^. Most intriguingly, a recently developed organoid-based 3D live-cell imaging assay demonstrated that breast cancer PDOs capture heterogeneous responses to T cells, intriguingly even within the same PDO culture^32^. This shows that PDO-based functional immunoassays can be used to screen for sensitivity to T-cell immunotherapies and, more importantly, to identify the subclonal cell populations that can potentially drive resistance. For ovarian cancer, short-term 3D cultures established from patient’s primary samples that preserve tumor-associated immune cells have captured a snapshot of the molecular changes in the NK compartment: an anti-tumoral molecular phenotype emerged after immune checkpoint inhibition, demonstrating the critical role of NK cells in therapeutic responses to immunotherapies and the potential of 3D *in vitro* systems to study NK cell activity in ovarian cancer^38^. All these findings encouraged us to explore whether variations in responsiveness to NK cells exist and could be captured in patient-derived ovarian cancer organoids.

Here, we report the development of an ovarian cancer PDO co-culture system with iNK cells in combination with 3D live-cell high content imaging technology. We found that responses to iNK cytotoxic activity vary between individual organoids within the same culture. We show that this heterogeneity was different between PDOs established from different ovarian cancer patients. In addition, we identified a rare subpopulation of organoids, present in both patient lines to different degrees, that are unresponsive to NK cell cytotoxicity. Our results provide evidence that ovarian cancer PDOs can be used to reveal subclonal response heterogeneity to NK cell-based immunotherapies and discern the differences between patients or tumor subtypes.

## Results

In this study we adapted published protocols^43,51–53^ to derive PDOs from a high grade serous ovarian carcinoma resected from the omentum (PDO-1) and an ovarian mucinous carcinoma resected from the left ovary (PDO-2) (Fig.1a). Immunohistochemical analysis revealed that PDOs expressed the ovarian cancer biomarker paired box gene 8 (PAX8), while tumor protein p53 expression was only detected in PDO-1 (Supplementary Fig.1). As a source of NK cells, we chose iNKs which we generated from iPSCs using established differentiation protocols^54–57^ (Fig. 1b). We consistently obtained >90% CD45+CD56+ cells that also expressed NK-specific inhibitory and activating receptors (Fig.1b-c) and displayed cytotoxic activity against the leukemia cell line K562 (Fig. 1d). To reduce line-to-line variability, we used only one iPSC line generated by reprogramming human hematopoietic stem/progenitor cells isolated from peripheral whole blood from a healthy donor. To further reduce assay variability, we generated a large batch of iNK cells that were then cryopreserved after phenotypic and functional validation. We also verified that cryopreserved iNKs maintain killing activity comparable to freshly differentiated iNKs in cytotoxic assays against K562 cells, confirming that they could be used in downstream killing assays (Fig. 1e). We next tested which media conditions enable simultaneous maintenance of ovarian cancer PDOs and iNKs in co-culture. First, we evaluated the viability of iNK cells exposed to organoid media (tOva) or a 1:1 (v/v) iNK-to-organoid media mixture (iNK:tOva media). We also tested viability in organoid media without nicotinamide (tOva-B3), a common component of PDO cultures. A previous study has shown that nicotinamide interferes with the functionality of CAR-engineered NK-92 cells in a patient-derived colorectal cancer organoid system^31^. When compared to iNK media, tOva and tOva-B3 conditions significantly decreased the amount of CD45+CD56+ cells after 24 h, although the remaining CD45+CD56+ cells remained viable. In contrast, iNK:tOva media conditions allowed the CD45+CD56+ population to remain stable and viable even after 24 h (Fig. 1f). To determine PDO viability in iNK:tOva and iNK media conditions, we performed cell viability assays using CellTiter Glow® 3D reagent to quantify ATP levels. Intact organoids were collected, counted using optimized pipelines with the Keyence hybrid cell count analysis software (Supplementary Fig. 2) and plated in 384-well plates at 1,000 organoids *per* well. In comparison to the viability in standard tOva media conditions, exposure to iNK media resulted in significant decrease in PDO-1 viability. In contrast, both PDO lines remained viable in iNK:tOva media for at least 20 h (Fig. 1g). Based on these results we chose iNK:tOva media conditions to develop our live cell imaging assay in iNK/ovarian PDO co-cultures.

**Figure 1.**
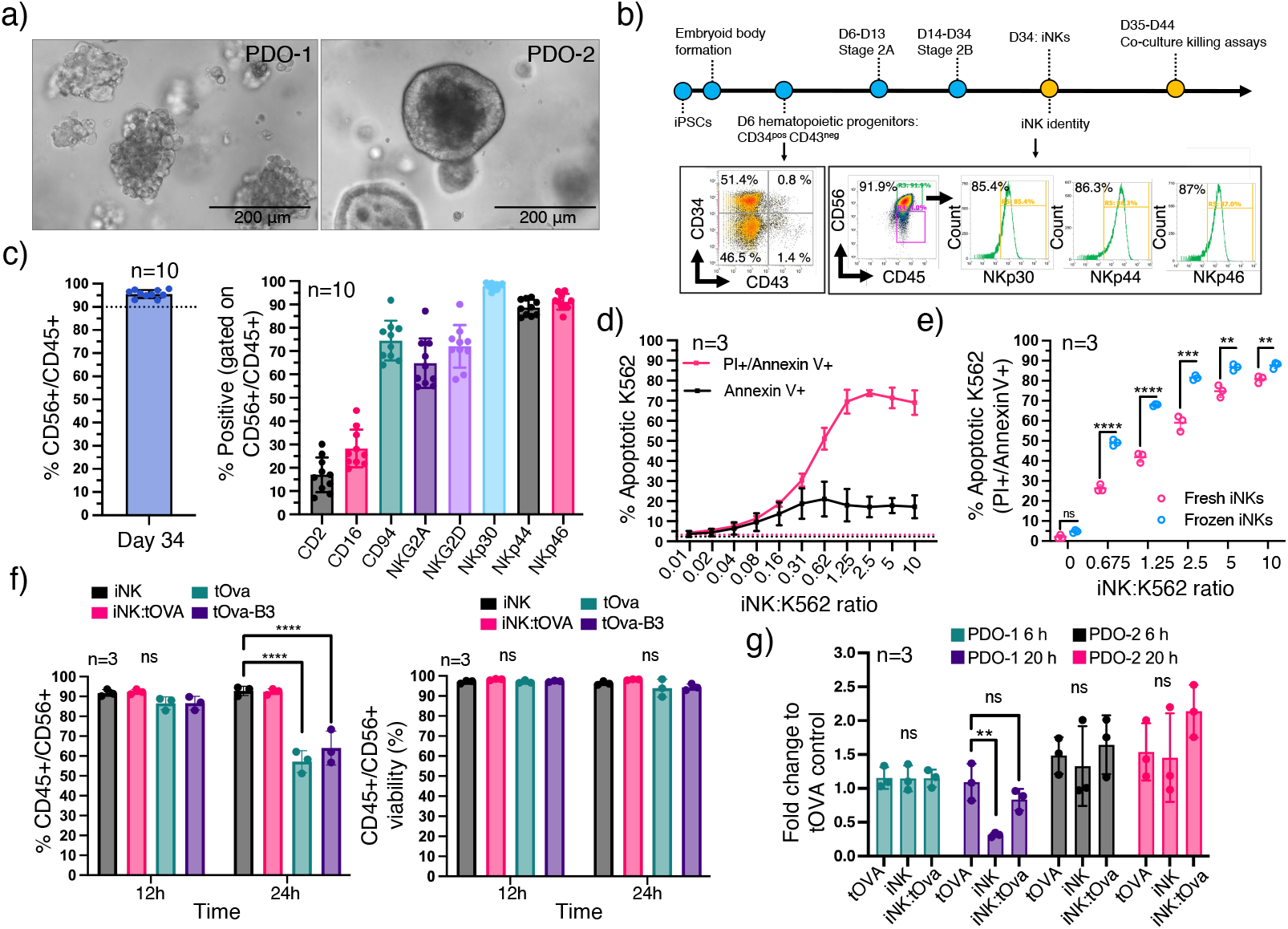
Patient derived ovarian cancer organoids and iPSC derived functional NK cells can be maintained in co-culture media conditions. (a) Representative bright-field images showing organoid morphology of PDO-1 (passage 8, p8) and PDO-2 (p3). Scale bar, 200 μm. (b) Schematic showing the differentiation protocol used to generate iNKs. Representative quality control analysis by flow cytometry to evaluate differentiation efficiency in each differentiation round; day 6, percentage of CD34^+^ hematopoietic progenitors and day 34, percentage of CD56^+^CD45^+^ (double positive) cells (iNK cells) from which expression of canonical NK cell surface markers are evaluated (representative flow cytometry plots for NKp30, NKp44, and NKp46 are shown). (c) Pooled flow cytometry data showing robust production of iNK cells (mean ±SD; n = 10 independent differentiation rounds from one iPSC line). (Left) percentage of CD56^+^CD45^+^ cells at day 34 after the initiation of cell differentiation. (Right) percentage of CD56^+^CD45^+^ cells expressing the indicated NK cell surface marker. Each dot represents one independent experiment. (d) Fraction of apoptotic cells in cytotoxic assays against K562 cells using a dilution series of effector (iNK) to target (K562) ratio (E:T) from 0.01:1 to 10:1 for 4h. Cell death was detected with Propidium Iodide and Annexin V or with Annexin V alone (mean ±SD; n = 3 independent experiments). (e) Fraction of apoptotic cells in cytotoxic assays against K562 cells comparing freshly differentiated iNKs to cryopreserved iNKs after overnight recovery. The E:T ratios are indicated. Cell death was detected with Propidium Iodide and Annexin V (mean ±SD; n = 3 independent experiments). (f) Survival of iNK cells in different media conditions at 12h and 24h in comparison to iNK media control. The plots represent pooled flow cytometry experiments to quantify the proportion of CD56^+^CD45^+^ cells (left) and the viability of the gated CD56^+^CD45^+^ population (mean ±SD; n = 3 independent experiments) (e, f) (Test for significant differrence: two-way ANOVA, Sidak’s multiple comparisons test, single pooled variance). (g) Pooled CellTiter Glow® 3D assays in PDOs showing their fold change under different media conditions as indicated at 6h and 20h in comparison to standard organoid media condition control (mean ±SD; n = 3 independent experiments) (significance test: two-tailed, unpaired t-test). *p<0.03; **p<0.002; ***p<0.0002; ****p<0.0001; ns = not significant

For live-cell imaging, formed organoids were collected intact from the Cultrex® Basement Membrane Extract (BME) they are embedded in at day 6 after plating, a timepoint at which most organoids range from 30-70 µm in diameter (Fig. 2a). The collected organoids were then labelled with CellTracker™ Orange CMRA, filtered using a 70 μm cell strainer and counted as described above (Supplementary Fig. 2). In parallel, iNKs were collected from an overnight culture after thaw and labeled with CellTrace™ Calcein Blue. Next, labeled iNKs and organoids were mixed in iNK:tOva media containing 5% Cultrex® BME, and plated in a pre-coated 96-well plate at 20:1 effector (iNKs) to target (organoids) (E:T) ratio *per* well^29,31^. To detect apoptotic events, CellEvent™ Caspase-3/7 Green detection reagent was added at the time of plating. This reagent consists of a peptide conjugated to a nucleic acid binding dye that only binds to DNA and produces a fluorogenic signal after cleavage by activated caspase-3/7 with reported signal stability up to 72 hours (h)^58^. As a control, labeled organoids were plated in the same conditions without iNKs in parallel wells (*i*.*e*. monoculture controls). Co-culture and monoculture controls were incubated for 30-45 minutes and then subjected to live cell imaging in an Opera Phenix™ High Content Screening system which utilizes an advanced confocal spinning-disk technology providing high 3D imaging resolution with four fluorescence channels imaged simultaneously^59^. Images were acquired every 15 minutes for up to 11h (Fig. 2b) capturing the tumor organoid-iNK cell interactions and caspase-3/7 activation dynamics across 500+ individual organoids simultaneously *per* condition (Fig. 2c). We next created an image analysis pipeline, using the Opera Phenix™ Harmony software, to quantify the area of each individual CMRA-labeled organoid positive for activated caspase-3/7 signal at each acquisition time point as a measurement of each individual organoid’s susceptibility to iNK cytotoxic activity (Fig. 2d-f, Supplementary Fig. 3). As evident from monoculture controls, our labeling and co-culture media conditions did not significantly induce apoptosis. In contrast, we observed a robust apoptotic signal in organoids co-cultured with iNKs that increased over time (Fig. 2e-f). We observed differences in response kinetics (Fig. 2e) as well as overall response (Fig. 2f) between the PDO lines studied. Importantly, individual organoids within the same PDO culture displayed varying levels of apoptotic signal suggesting that tumor immune heterogeneity is preserved in ovarian cancer PDOs (Fig. 2f). Interestingly, we also observed that, in both PDO lines, a small fraction of organoids displayed no apoptotic events throughout the assay which we hypothesize could correspond to resistant subpopulations (Fig. 2g).

**Figure 2.**
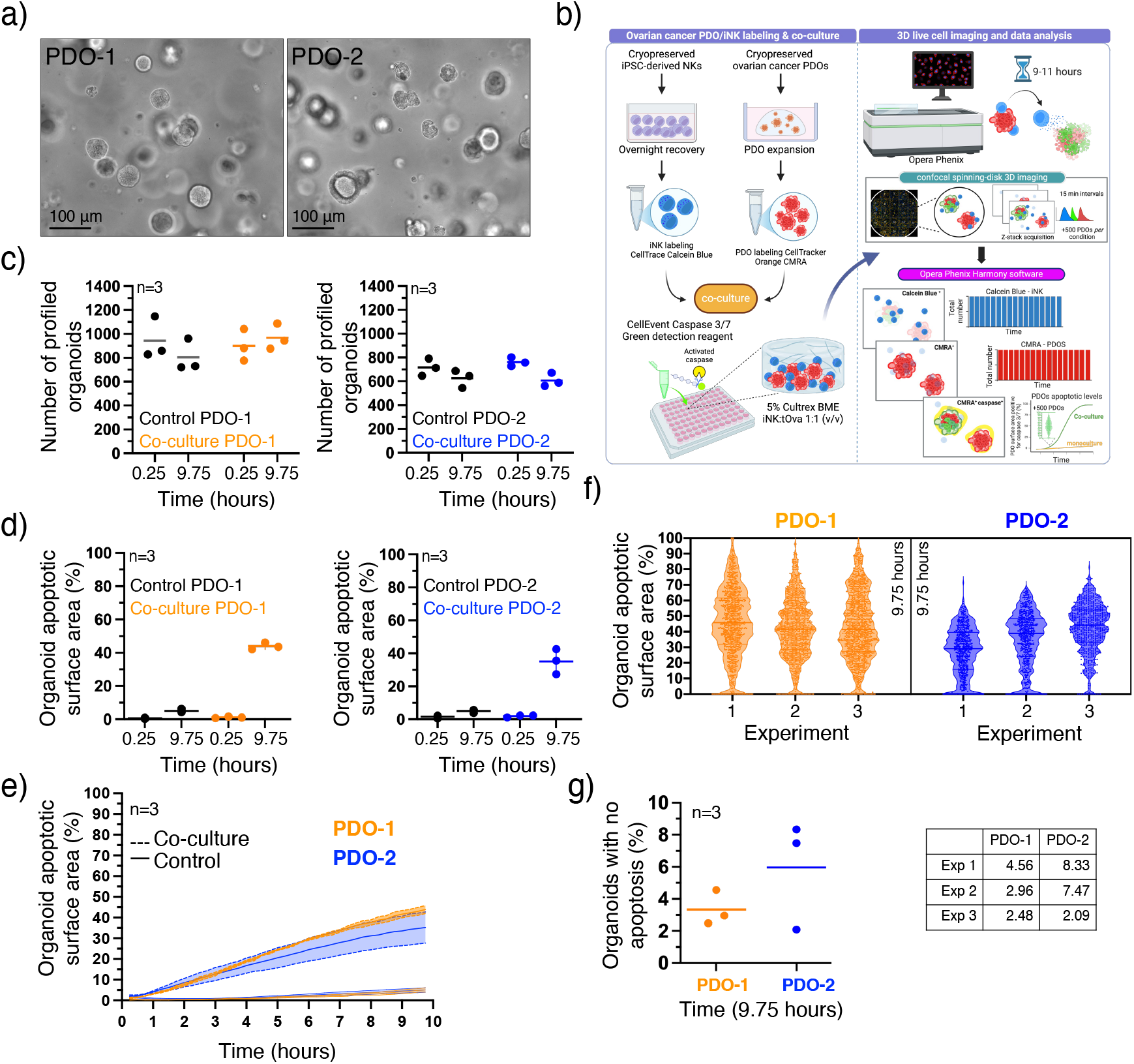
Inter- and intratumor response heterogeneity to iNKs cytotoxic activity. (a) Representative bright field images showing PDO-1 and PDO-2 cultures at the time of collection for co-culture experiments. Scale bar, 100 μm (b) Schematic representation of the 3D live cell imaging killing assay in PDO/iNK co-cultures (c) Total number of individual organoids captured per experiment and (d) average organoid area positive for caspase-3/7 in each individual organoid *per* condition (monoculture control *vs*. co-culture) in PDO-1 and PDO-2. The start- and endpoint of the assay are shown. Each data point represents one independent experiment [(c) mean; (d) mean ± SD]. (e) time course capture (15 minutes intervals) and organoid death quantification from 3D live cell imaging assays in PDO-1 and PDO-2 in monoculture controls *vs*. co-cultures (pooled 3 independent experiments; mean ± SD). (f) violin plots of apoptotic area in individual organoids, showing heterogeneous susceptibility to iNKs in PDO-1 and PDO-2 after 9.75h in co-culture. (g) fraction of organoids with no apoptotic events at the end of the assay (mean). Each data point represents one independent experiment. The specific percentage is shown in the table.

We next sought to further characterize the resistant and sensitive organoid subpopulations and compare their frequency between the studied PDO lines. Thus, we applied a Bayesian Beta mixture model with two components to the distribution of organoid apoptotic area at the experimental endpoint (Fig. 3a). With this model we inferred that the fraction of sensitive organoids was 0.93 (CI:0.92-0.94) and their average apoptotic area was 43.8% (CI:43.3-44.5%) whereas the resistant population had an average apoptotic area of 12.0% (CI:10.1-14.6%). Based on the modeling results, we found a threshold of 8.6% apoptotic area area to distinguish optimally between resistant and sensitive organoids. By applying this threshold across all time points of the experiment, we found that the mean apoptotic area increased from 13.5-23.1% at the initial quantification time point to 32.0-49.1% during the experiment but was stable for non-apoptotic organoids, resembling the control range (Fig. 3b). Moreover, we found that the proportion of apoptotic organoids increased sharply at the beginning of the experiment and thus, by 5 h the majority of organoids (>75%) displayed apoptotic events in both PDO co-cultures (Fig.3c). Towards the end of the experiment the proportion of apoptotic organoids plateaued in a sigmoid fashion, indicating a persisting population of resistant organoids. In contrast, most organoids (>90%) remained alive in monoculture controls throughout the duration of the assay (Fig. 3c). We then tested logistic, Hill and Gompertz models for fitting the kinetics of organoid apoptosis and found the Gompertz model to fit best to both datasets (*p*<0.001 and *p*<0.004 respectively) (Supplementary Fig.4). From this model we estimated, for both PDO lines combined, a resistant organoid fraction of 6.2% (CI:4.9-7.5%) and a difference of 1.4% (CI:0.2-2.8%) in resistant fraction between PDOs (Fig. 3d). Intriguingly, we were able to visualize iNKs actively engaging with organoids with low or entirely absent apoptotic signals throughout the duration of the assay (Fig. 4). We next evaluated if individual organoid size could be a predictor of the observed differential response. When we looked at the relationship between organoid diameter and apoptotic area in the monoculture controls, it showed a weak negative relationship indicative of smaller organoids exhibiting higher apoptotic area rates with Pearson’s correlation (R) ranging from -0.084 to -0.25 across replicates (Fig. 3e). Interestingly, in the co-culture conditions, we observed two subpopulations at the endpoint (31% are small organoids with a mean of 28 µm diameter and 69% are larger ones with a mean of 247 µm diameter) (Supplementary Fig. 5) with resistant organoids enriched in the small subpopulation (odds ratio=3.64, *p*<2e-16, Fisher’s exact test), for both PDO lines (Fig.3f). Sensitive organoids were found in both the small and large subpopulation. This might indicate that small size is necessary but not sufficient for a resistant response. Intriguingly, PDO lines differed in the relationship between size and apoptotic area: While sensitive organoids in PDO-1 showed a negative relationship between organoid diameter and apoptotic area across the three replicates, this relationship was positive in PDO-2 (*R*=0.43-0.59) (Fig. 3f).

**Figure 3.**
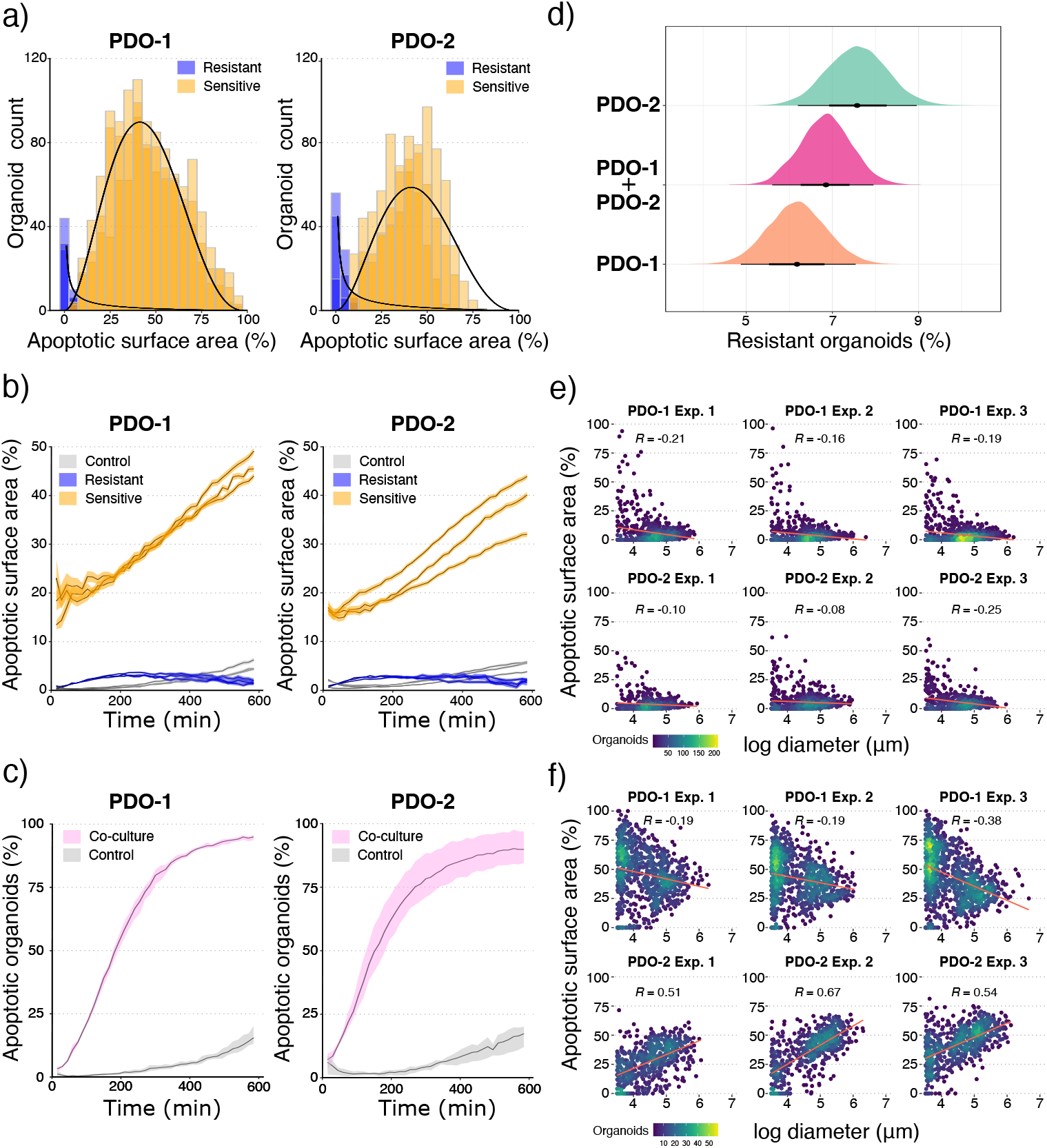
Ovarian cancer PDO cultures contain organoids resistant to iNK cytotoxic activity. (a) Organoid apoptotic area in PDO-1 and PDO-2 at the final time point. Density curves for a fitted beta mixture model are shown, where the two components correspond to resistant organoids (small apoptotic area, blue) and sensitive organoids (large apoptotic area, orange). Three independent experiments are shown for each PDO culture depicted as different color intensity. (b) Mean apoptotic area over time in PDO-1 and PDO-2. iNK-treated organoids are split into resistant (blue) and sensitive groups (orange). Monoculture controls are also shown (grey). Three independent experiments are shown for each PDO culture. Error bands correspond to the mean ± 2 SEM across all individual organoids quantified in each time point). (c) Fraction of apoptotic organoids (orange) over time in PDO-1 and PDO-2. Changes in monoculture control are also shown (light grey); Error bands represent the minimum and maximum percentage across three independent experiments. (b,c) Organoid populations (sensitive, resistant) are defined based on a threshold of apoptotic area inferred using a Bayesian beta mixture model. (d) Density plots of the percentage of resistant organoids in each PDO inferred using a Bayesian Gompertz model. The PDO-1+PDO-2 group indicates the values for a model without a coefficient for the patient, where the groups are merged. The median and the 95% and 80% credible intervals are shown by the dot, bold bar and thin bar respectively. (e-f) Organoid apoptotic area versus log diameter across PDOs in three independent experiments in monoculture controls (e) and the iNK co-culture condition (f) at the final time point. Color scale in scatter points indicates point density. The Pearson’s correlation (R) for each plot is shown.

**Figure 4.**
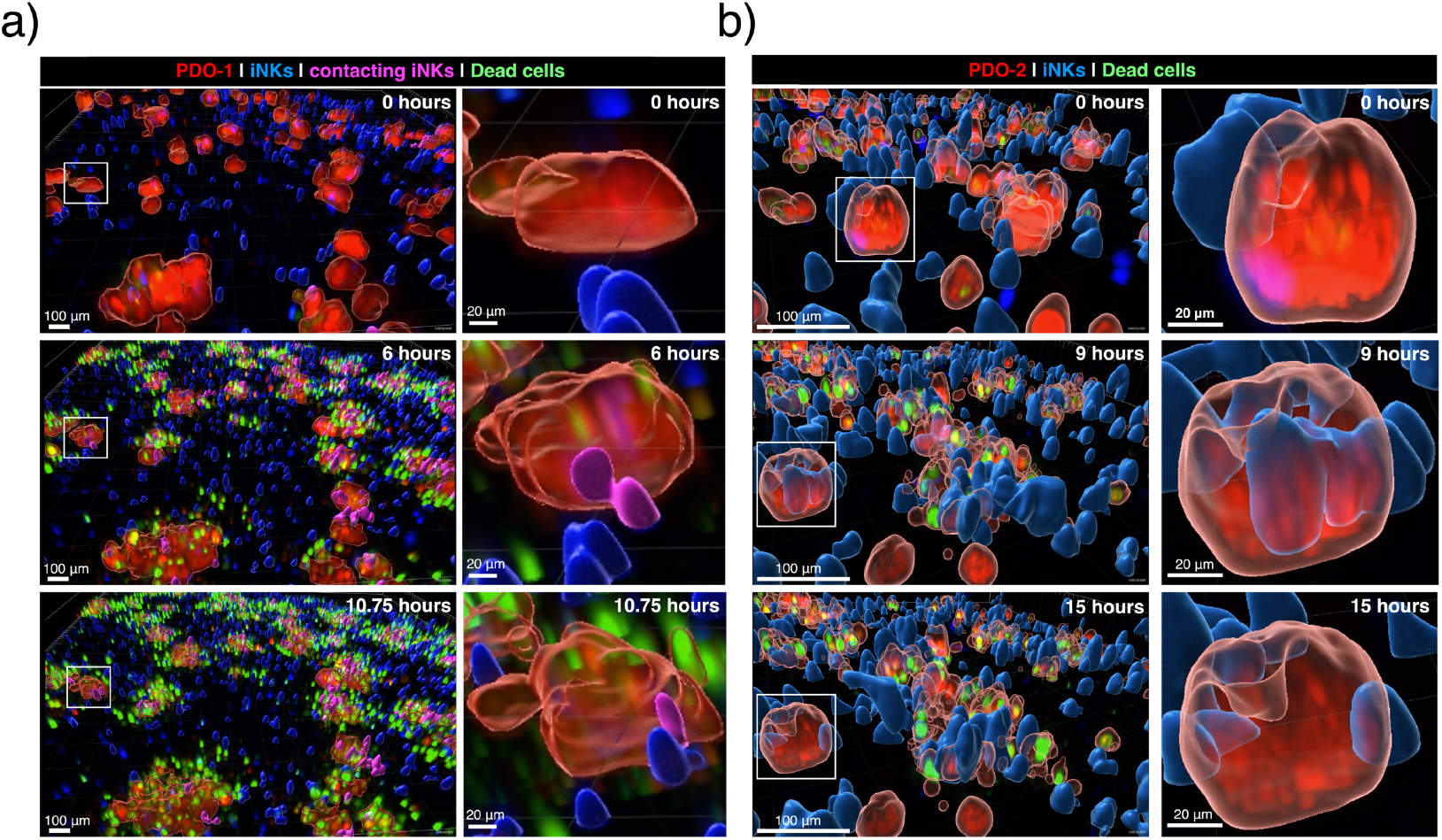
Intra-tumor organoids resistant to iNK cytotoxic activity. Processed images 3D-rendered with Imaris software (Oxford instruments) showing (a) PDO-1 and (b) PDO-2 organoids in co-culture with iNKs at different time intervals during one representative live cell imaging assay experiment. Images were acquired in Opera Phenix™ at (a) 10x or (b) 20X for increased resolution of tumor cells and iNK interactions. (a) Cells colored in magenta correspond to iNKs contacting with organoids defined as those located at a 0 μm distance from the corresponding organoid (Imaris software). The white square in the left panels, frames the organoid presented in the right panels showing an iNK-resistant organoid in each PDO co-culture. Scale bars, 100 μm (left panels) and 20 μm (right panels).

## Discussion

PDOs in co-culture with immune cells are emerging as powerful *in vitro* models to assess and optimize immunotherapeutic strategies for treating solid tumors. A variety of systems have been established but for the most part these capture organoid culture responses in bulk precluding the observation of sub-clonal response differences to immune cells that may be recapitulated in organoid culture systems. A novel 3D imaging and analysis technique developed by Dekkers *et al*^32^, revealed functional heterogeneity of engineered T cells in co-culture with breast cancer PDOs. Intriguingly, this system also revealed inter- and intra-PDO response heterogeneity, suggesting that PDOs also capture the heterogeneous responses to T cells. Therefore, we sought to develop a system for ovarian cancer as the high level of heterogeneity presented in this cancer drives frequent relapses with tumors resistant to treatments. However, T-cell based immunotherapies present several challenges to treat ovarian cancer which has led to limited efficacy^60^. In contrast, emerging evidence strongly supports the development of NK cell-based immunotherapies to treat ovarian cancer^18,21^. Here, we present an ovarian cancer PDO co-culture system with iNK cells and 3D live-cell imaging assays to quantify, in real time, iNK-mediated apoptotic cell death across hundreds of individual organoids. Using this assay we have reproducibly captured a spectrum of responses to iNK cell-mediated cytotoxic activity (Fig. 2). Among these we have identified a rare subset of individual organoids that remain intact even after active engagement by iNK cells, suggesting that these organoids potentially represent a resistant sub-clonal population (Figs. 3-4).

NK cells overcome key limitations of T cell-based therapies, such as cytotoxicity independent of antigen presentation or HLA expression, and feasibility for allogeneic use. Thus, adoptive NK cell transfer therapy including NK cells derived from iPSCs, are among the most attractive options as NK cells do not induce graft-versus-host complications in the allogeneic setting presenting an unprecedented potential as off-the-shelf therapy^61^. For ovarian cancer, positive results have been reported in clinical trials testing NK cell-based therapy, including heavily pre-treated patients^19^. NK cells have also been shown to be overrepresented in the ovarian tumor immune infiltrate in comparison to circulating blood cells^20,62^, are effective at killing ovarian cancer cells^63^, and a higher percentage of NK cell infiltrate correlates with an overall survival benefit^17,22,64,65^. However, the detailed mechanisms that drive responses to NKs in ovarian tumors remain largely unexplored^20,63,66^. Given the complexity and high level of heterogeneity displayed by ovarian cancer tumors, understanding the varying interactions of ovarian cancer cells with NK cells in each patient will enable effectively harnessing their potential to treat ovarian cancer. Previously, 3D co-culture systems have been developed to evaluate responses to NK cell-based therapies for the most part using cancer cell line spheroids^27,28,31,67–72^. For ovarian cancer, short term organoid cultures that maintain immune cell populations, including NK cells, have been used to characterize changes in these populations after immune checkpoint inhibition^38,73^. But these systems lack expansion capabilities, and the scalability for testing multiple experimental conditions. Additional NK co-culture systems have mainly relied on ovarian cancer cell line spheroids^28,29^, lacking the complexity offered by PDOs established from patient-derived material. Our assay is robust, reproducible and able to quantify the effects of NK cell-mediated apoptotic events in real time across hundreds of individual organoids and between different PDO cultures. Our assay provides the means to correlate responses with tumor-iNK cell interactions and captures resistant and sensitive phenotypes within the same tumor, allowing assessment of the potential effectiveness of NK cell-based therapies in candidate patient populations. Intriguingly, our system captures individual organoid responses including those that may drive therapy resistance and is amenable to functional immunotherapy and/or compound testing, thereby opening avenues with which to study patient specific responses and optimize NK-cell based therapies for ovarian cancer. Our technology offers a promising platform for functional testing that can be adapted to other solid tumor types and/or immune cells so that the effects of immune modulatory agents can be assessed and optimized to enhance immune cell killing activity in resistant cells.

## Supplementary Figures

**Supplementary Figure 1.**
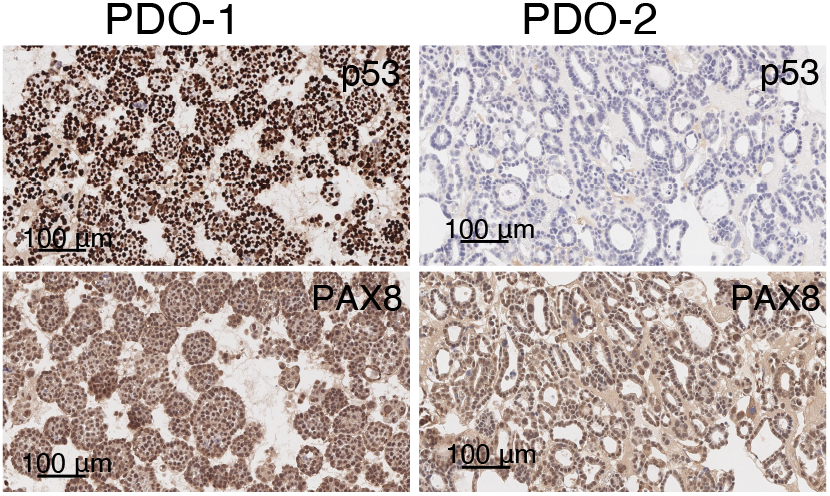
p53 and PAX8 phenotype in ovarian cancer PDOs. Representative p53 and PAX8 immunohistochemistry in PDO-1 and PDO-2. Scale bars, 100 μm.

**Supplementary Figure 2.**
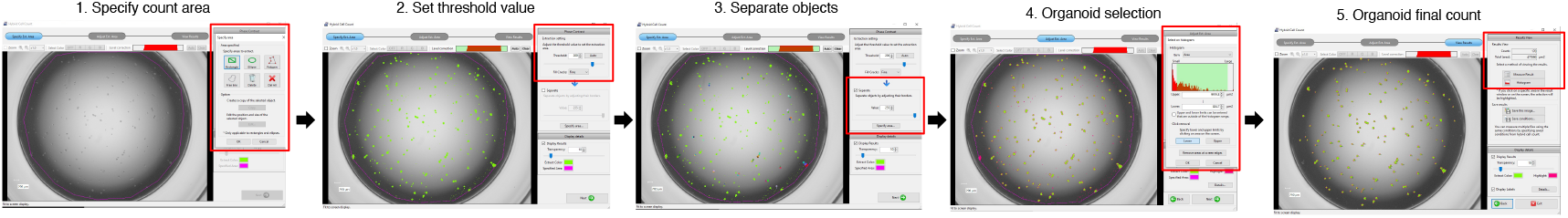
Organoid count pipeline. Organoid suspensions were counted using the Keyence hybrid cell count analysis software to determine their concentration for precise plating. For counting, an aliquot of the organoid suspension was plated in 96-well plates in triplicate and phase contrast images were taken with a Keyence BZ-X810 microscope for analysis. (a) Representative screenshots showing the counting organoid pipeline using Keyence hybrid cell count analysis software: (1) the count area is defined excluding the edges and the exterior of the well; (2) using the extraction setting, the threshold value for the objects that will be counted is set; some cellular is excluded at a later step; (3) organoids are cunted that have aggregated as individual organoids, objects are separated by adjusting their borders; (4) the organoids to be counted are then selected by gating them in an histogram of organoid size. Masked objects corresponding to debris and aggregated organoids are excluded; (5) organoid count is obtained. The count average across the three replicates is considered as the final count.

**Supplementary Figure 3.**
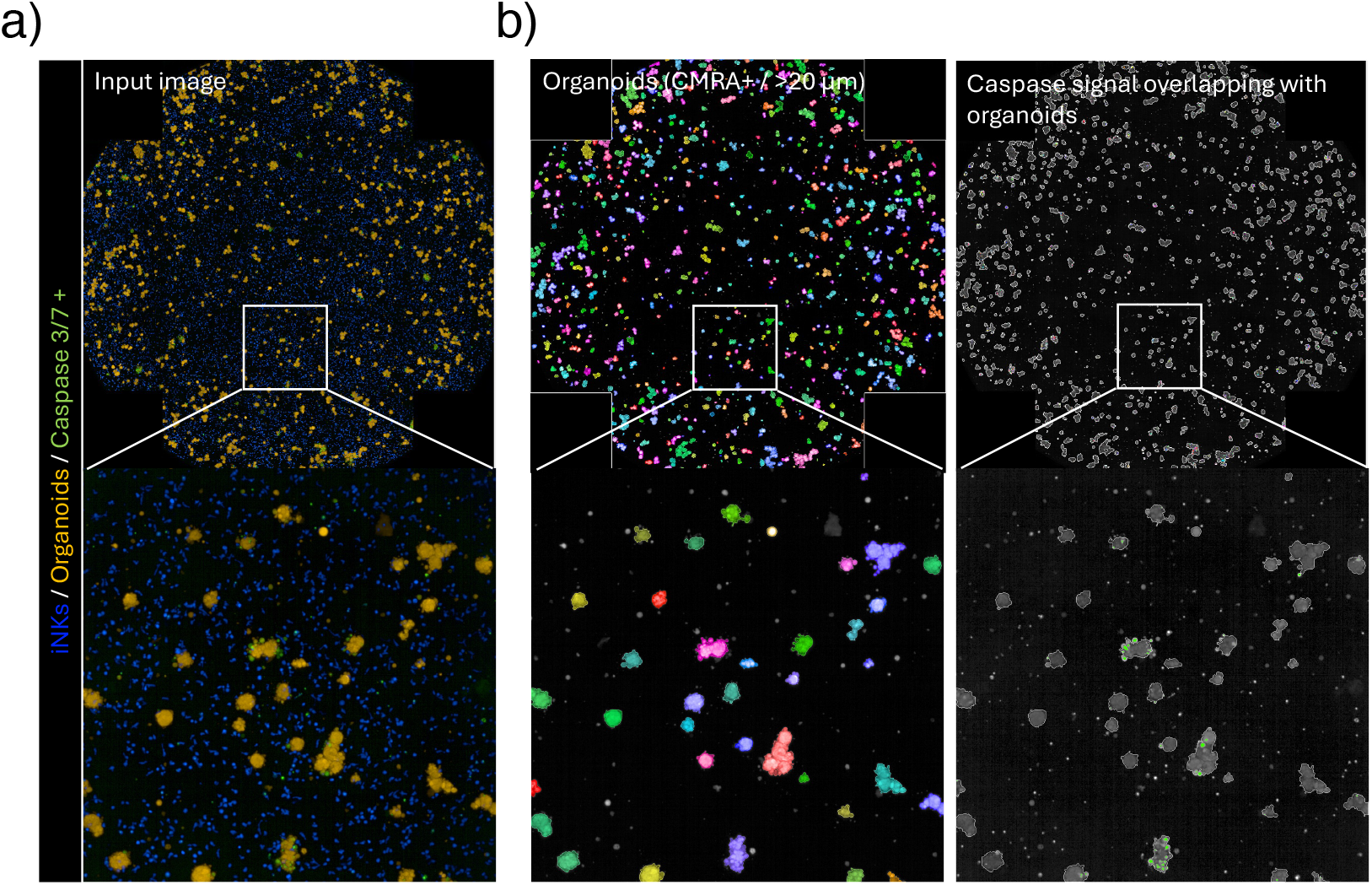
Identification of Caspase 3/7 positive signals in PDOs in co-culture conditions. (a) Representative images acquired with Opera Phenix™ High Content Screening system showing PDO-1 (CMRA positive, orange) in co-culture with iNKs (Calcein Blue positive, blue) in the presence of caspase-3/7 reagent (green signals) at the starting of the assay. The top image corresponds to the entire well. The bottom image shows a magnification from the inlet. (b) Corresponding identification for quantification of (left) organoids (CMRA positive objects that are >20 μm) and (right) overlapping caspase-3/7 signals with Opera Phenix™ Harmony software. The colors (left) and contour (right) in organoids correspond to the mask assigned by the software.

**Supplementary Figure 4.**
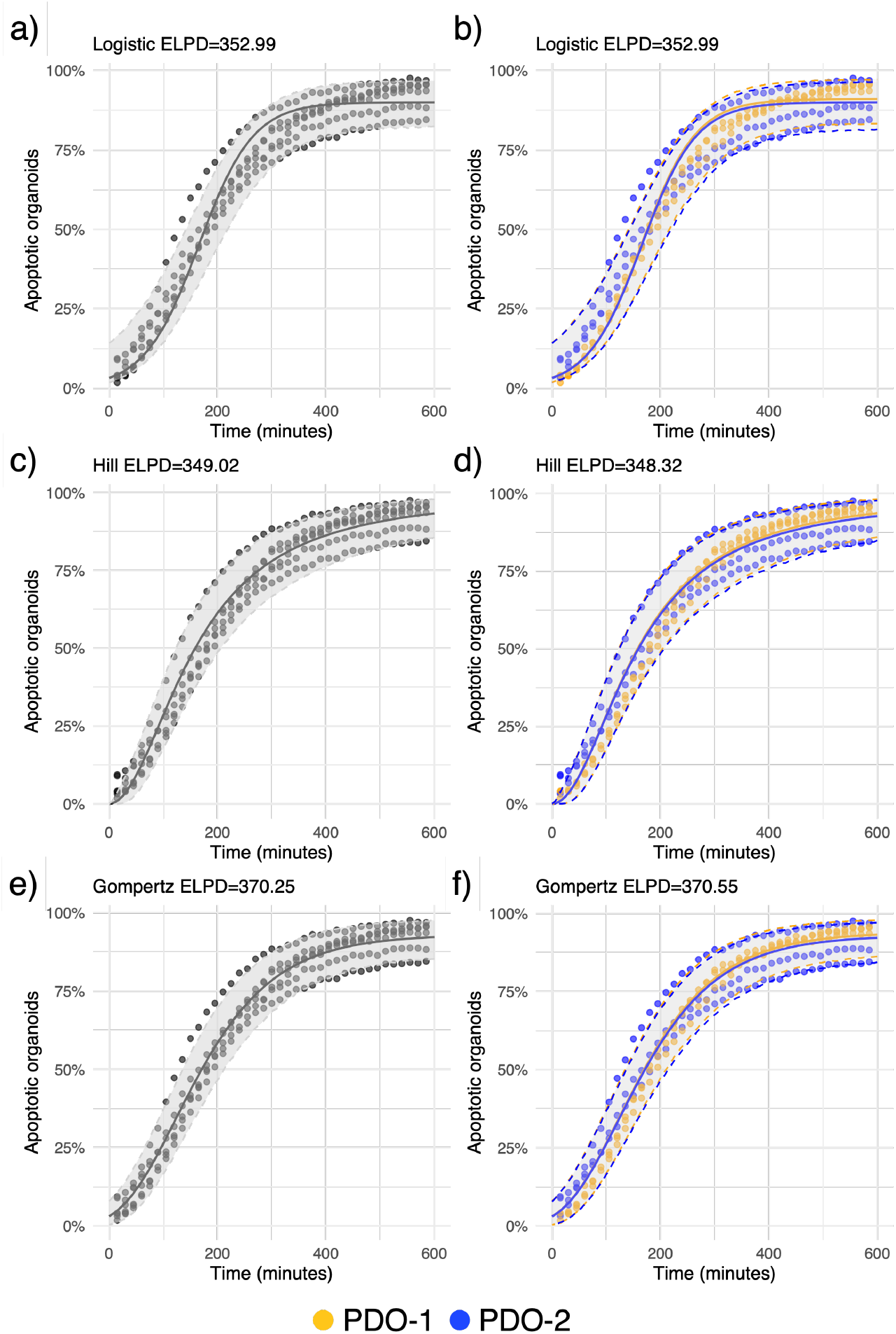
Comparison of sigmoid fitting to percentage of apoptotic organoids over time. A logistic model fit without (a) or with (b) an organoid patient coefficient is compared to Hill (c and d) and Gompertz (e and f) models. The expected log predictive density (ELPD) is shown for each model, with greater numbers indicating better fit. The 95% credible intervals for each curve are shown.

**Supplementary Figure 5.**
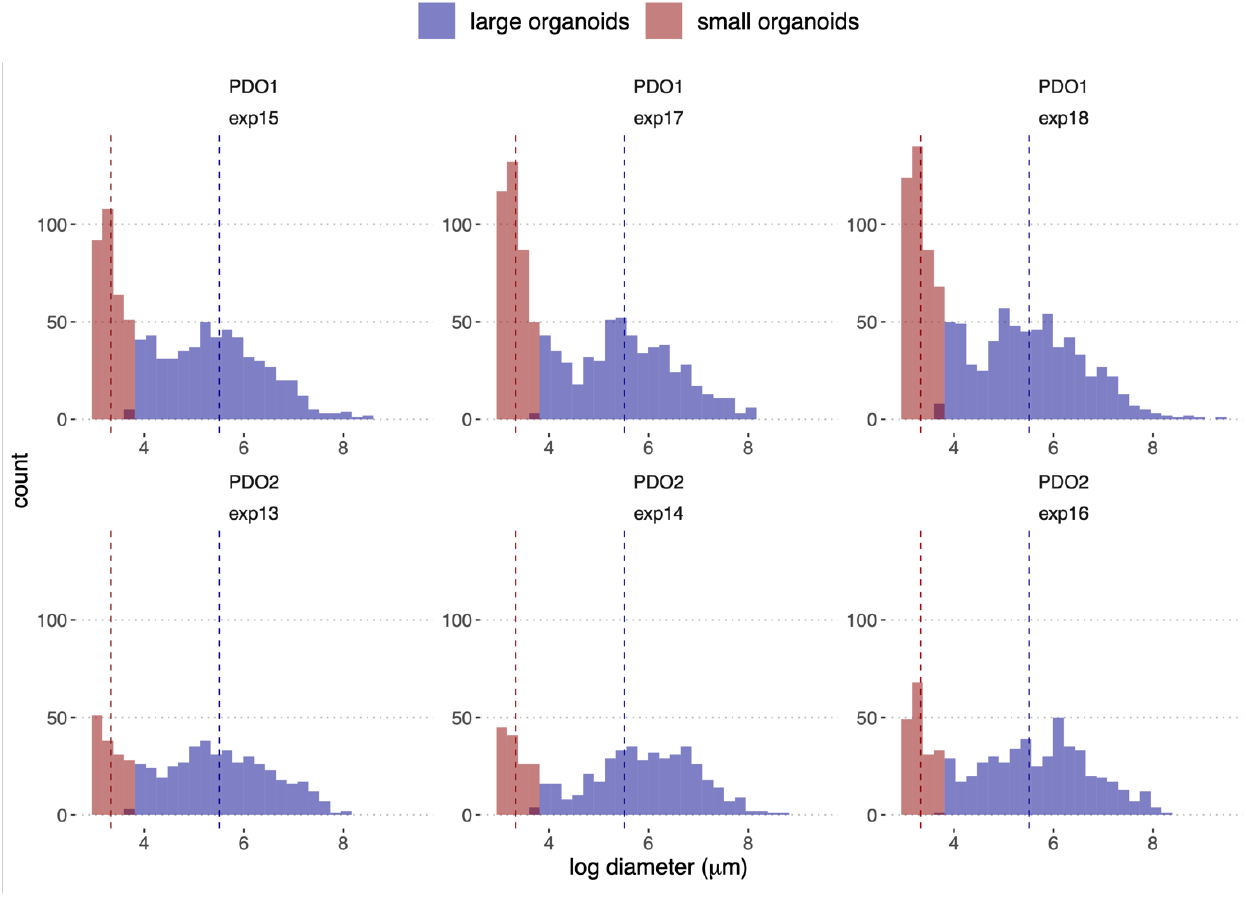
Organoid size distribution across PDOs in the co-culture condition at the endpoint of the assay. Three independent experiments for each PDO culture are shown. Organoids are assigned to the group of small or large organoids based on a Gaussian mixture model. Fitted means of the small and large organoid groups are indicated with vertical dotted lines.

## Acknowledgements

We would like to thank the patients and families that consented to donate the biospecimens that were used to establish the PDOs. We thank Matt Zimmer and Gist Croft for technical input. Howard Kim, Cecile Terrenoire, Fawaz Saleh, and Alex Annealing for kindly providing the iPSC line used in this study and for technical support. We thank Dr. Dmitriy Zamarin from Memorial Sloan Kettering Cancer Center (MSKCC) for providing tumor samples. Figure 2b was created with Biorender.com. This work was supported by the NCI R21CA240219 (LM), the New York Stem Cell Foundation Research Institute (NYSCF), and The Ralph and Ricky Lauren Family Foundation.

## Author contributions

Conceptualization: LM and YW. Tumor sample processing and organoid derivation: JH. Organoid culture and biobank: MM, TV, WS. iNK production, culture, characterization, and biobanking: YW, JE, MM. iNK K562 functional assays: YW, JE, NH. Co-culture optimization: MM, JE. Organoid collection, quantification, and labeling: WS, JSC, MM. iNK labeling: YW and JE. Co-culture and live cell imaging: JE, WS, JSC, MM. Quantification pipelines: LM, YW, TR, MS, JE, JSC, WS. Data analysis: AS, LM, JE, MM, SS, YW. Supervision: LM, YW, SS, FJM. Resources and funding acquisition: LM, FJM. Writing original draft: LM, AS, YW. Writing-review and editing: LM, YW, SS, FJM, AS, JSC, WS, MM.

## Declaration of interest

### Lead contact

For inquiries regarding the availability of materials and resources, contact Laura Andres-Martin, Yinan Wang, and Stefan Semrau.

### Materials availability

NYSCF Research Institute organoid models and associated clinical data, iPSC lines and iNKs may be made available on request through the NYSCF Repository (https://nyscf.org/partnering/products/), upon material transfer agreement.

## Experimental procedures

### Patient’s samples

De-identified tumor material was obtained from female patients diagnosed with ovarian cancer. Biospecimens were provided by MSKCC from consented patients and the Cooperative Human Tissue Network (CHTN), which is funded by the National Cancer Institute. Other investigators may have received specimens from the same subjects.

### Patient-derived ovarian cancer organoids derivation and culture

Patient-derived organoids (PDOs) were established from fresh tumor resections as described previously^43,53^ with slight modifications. Briefly, tumor tissues were digested in Gibco™ Advanced DMEM/F12 media (ThermoFisher Scientific, Cat. No. 12634-028) containing 1x Gibco™GlutaMAX (ThermoFisher Scientific, 35050-079), 1x Penicillin Streptomycin (10,000 U/ml) (Life Technologies, Cat. No. 15140122), 10 mM HEPES (ThermoFisher Scientific, 15-630-080), 100 µg/mL Primocin® (InvivoGen, ant-pm-1), 10 µm Y-27632 ROCK inhibitor (AbMole, cat. No. M1817) and 0.7 mg/ml Collagenase XI (Sigma C9407) for 25 minutes at 37ºC. The digested tissue was then filtered through a 100 µm cell strainer and centrifuged at 300g for 5 minutes at 4ºC. When needed, the resulting pellet was treated with 3mL of Red Blood cell Lysis buffer (Sigma, Cat. No. 11814389001) for 5 minutes at room temperature and then centrifuged at 300g, 4ºC for 5 minutes. For plating, the pellet was resuspended in an ice-cold 100% Cultrex® Reduced Growth Factor BME, Type 2, PathClear™ (Cultrex® BME) (R&D Systems, Cat. No. 3533-010-02) and droplets containing 4000-10,000 cells/µl were plated on pre-warmed multiwell tissue culture plates (Greiner CELLSTAR®, Cat. No. 677102). After plating, plates were placed in the incubator a 37ºC to allow the Cultrex® BME to solidify for 30-60 minutes a 37ºC. Once solidified, the domes with embedded cells were topped with pre-warmed ovarian tumor organoid medium (tOva) composed of Gibco™ Advanced DMEM/F12 (ThermoFisher Scientific, Cat. No. 12634-028), 1x Gibco™GlutaMAX (ThermoFisher Scientific, 35050-079), 1x Penicillin Streptomycin (10,000 U/ml) (Life Technologies, Cat. No. 15140122), 10 mM HEPES (ThermoFisher Scientific, 15-630-080), 100 µg/mL Primocin® (InvivoGen, ant-pm-1), 1x Gibco™ B-27™ Supplement (50x), (Life Technologies, Cat. No. 17504-044), 1.25 mM N-Acetyl-L-cysteine (Sigma-Aldrich, Cat. No. A9165), 10 mM Nicotinamide (Sigma-Aldrich, Cat. No. N0636), 0.5 µM A83-01 (Tocris, Cat.No. 2939), 0.5 µg/mL Hydrocortisone (Sigma-Aldrich, Cat. No. H0888), 10 µM Forskolin (R&D Systems, Cat.No.1099), 100nM β-Estradiol (Sigma-Aldrich, Cat. No. E2758), 16.3 µg/ml Bovine Pituitary Extract (BPE) ThermoFisher Scientific, Cat. No. 13028014), 10 ng/mL Recombinant Human FGF-10 (PeproTech, Cat. No. 100-26), 5 ng/mL Recombinant Human KGF (FGF-7) (PeproTech, Cat. No. 100-19), 37.5 ng/mL Recombinant Human Heregulin Beta-1 (PeproTech, Cat. No. 100-03), 5 ng/mL Recombinant Human EGF (PeproTech, Cat. No. AF-100-15), 100ng/ml

Recombinant Human R-Spondin-1 (PeproTech, 120-38), 1% Noggin-Fc Fusion Protein Conditioned Medium (Immunoprecise, Cat. No. N002), 0.5 nM WNT Surrogate-Fc fusion protein (ImmunoPrecise, Cat. No. N001) and containing 10 µm Y-27632 ROCK inhibitor (AbMole, cat. No. M1817) and then incubated for organoid formation with media changes every 2–3 days and passaging every 1-2 weeks for expansion and biobanking. For biobanking, dense organoid cultures were cryopreserved as small cell clusters/single cells in Recovery™ Cell Culture Freezing Medium (Thermo Fisher Scientific, Cat. No. 12648010) at a concentration of 0.8-1.2×10^6^ cells *per* 500 µl. For co-culture and live cell imaging experiments, ovarian cancer PDOs were retrieved from the NYSCF organoid biobank. Cryopreserved organoids were thawed, plated and expanded for at least 7 days prior to use in co-culture experiments. Briefly, after thawing, cells were resuspended in 70-100% (depending on the specific PDO line requirements) cold Cultrex® Reduced Growth Factor BME, Type 2, PathClear™ (Cultrex® BME) (R&D Systems, Cat. No. 3533-010-02) and droplets were plated on pre-warmed multiwell tissue culture plates (Greiner CELLSTAR®, Cat. No. 677102). Droplets were allowed to solidify at 37°C inside an incubator for at least 30 min., then overlaid with tOva medium as described above containing 10 µm Y-27632 ROCK inhibitor (AbMole, cat. No. M1817). The medium was changed every 2–3 days for 7–10 days until dense organoid cultures are formed with organoids ranging in size from 100 to 500 µm. Organoids were then passaged with Gibco™ TrypLE™ Select Enzyme (ThermoFisher Scientific, Cat. No. 12563011), containing 10 µm Y-27632 ROCK inhibitor (AbMole, cat. No. M1817) and 10 µg/mL DNase I (Sigma-Aldrich, Cat. No. DN25), resuspended in 70-100% cold Cultrex® Reduced Growth Factor BME, Type 2, PathClear™ (Cultrex BME) (R&D Systems, Cat. No. 3533-010-02) plated in droplets at 800 cells/µL and cultured for further expansion or use in co-culture experiments.

### iPSC generation and culture

Peripheral whole blood was obtained from Comprehensive Cell Solutions, an operating division of the New York Blood Center, Inc. Following Ficoll separation, PBMCs were sorted to obtain hematopoietic stem/ progenitor cells (EasySep™ Human Progenitor Cell Enrichment Kit, Stemcell Technologies, Catalog #17936,) and cultured for three days in media containing IL-3 (CellGenix, Cat. No. 1002-050), IL-6 (CellGenix, Cat. No. 1004-050), FLT-3 (CellGenix, Cat. No. 1015-050), TPO (CellGenix, Cat. No. 1017-050), and SCF (CellGenix, Cat. No. 1018-050) to enrich for CD34+ cell population, after which they were reprogrammed using CytoTune™-iPS 2.0 Sendai Reprogramming Kit (Thermo Fisher, Catalog # A16517) according to manufacturer instructions. Individual clones were manually picked and passaged/expanded in Gibco™ Essential 8™ Medium (Thermo Fisher Scientific, Catalog # A1517001) on Recombinant human truncated Vitronectin-coated tissue culture-treated vessels (Gibco™ CTS™ VTN-N, Thermo Fisher Scientific, Catalog # CTS279S3) using TrypLE (Gibco™ CTS™ TrypLE™ Select Enzyme, Thermo Fisher Scientific, Catalog # A4738001), and cryopreserved in BiolifeSolutions® CryoStor® CS10 Cell Freezing Medium according to the manufacturer instructions (StemCell Technologies, Cat. No. 07930).

### Natural killer cells generation from iPSCs and culture

iPSCs were retrieved from our biobank and expanded as described prior to their differentiation into Natural Killer cells (iNKs). For iNK generation we used a spin embryoid body (EB) protocol adapted from previously published procedures^1,2^. For EBs formation, iPSCs were treated with CTS™ TrypLE™ Select (Thermo Fisher Scientific, A1285901) to yield a single cell iPSC suspension. To render one EB per well, iPSCs were seeded into an entire 96-well Clear Round Bottom Not-TC Treated Microplates (Falcon®, Corning, 351177) at 8,000 per well, in 150 µL of STEMdiff™ APEL™2 Medium (StemCell Technologies, Cat no 05270) containing 50 ng/mL recombinant human bone morphogenetic protein 4 BMP-4 (R&D Systems, Cat no 314-BP/CF), 20 ng/mL recombinant human FGF basic (bFGF) (146 aa) Protein (R&D Systems, Cat. No 233-FB), 50 ng/mL recombinant human vascular endothelial growth factor (VEGF) 165 Protein, CF (R&D Systems, Cat. No 293-VE), 50 ng/mL recombinant human stem cell factor SCF (Peprotech, Cat. No 300-07) and 10 µM Y-27532, ROCK inhibitor Y-27532 (StemCell Technologies, 72302).

After spinning the plates at 300 xg for 5 min at room temperature (RT), they were cultured at normoxic (20% O2, 5% CO2) condition in a 37°C incubator for 6 days. After 6 days, 15 EBs were collected and plated into one well of a 6-well TC-treated culture vessels coated with 10 µg/mL laminin 521 (BioLamina, MX521). For coating, plates were incubated for 1 hour at 37 °C. EB were then cultured for 28-30 days (D28-D30) in 5 mL NK differentiation medium with half volume media changes every 2-4 days. Using this protocol approximately 1–5×10^6^ iNK cells can be collected from one single 6 well. During the first 7 days, the NK differentiation medium consisted in 85% 2:1 (v/v) mixture of Gibco™ DMEM (ThermoFisher Scientific, Cat. No. 10569010) and Gibco™ Ham’s F-12 Nutrient Mix (ThermoFisher Scientific, Cat. No. 31765035), 15% human AB serum (GeminiBio, Cat. No. 100-612), 1x Gibco™ MEM Non-essential amino acids (NEAA) (Thermo Fisher Scientific, Cat. No. 11140050), 1x Gibco™ GlutaMAX™ Supplement (Thermo Fisher Scientific, Cat. No. 35050061), 1x Gibco™ Penicillin/Streptomycin (50 U/mL) (ThermoFisher Scientific, Cat. No. 15070063) 25 µM Gibco™ 2-Mercaptoethanol (Thermo Fisher Scientific, Cat. No. 21985-023), 5 ng/ml Sodium selenite (Sigma-Aldrich, Cat. No. S5261), 50 µM Ethanolamine (Sigma-Aldrich, Cat. No. E0135), 20 mg/L L-Ascorbic acid (Millipore Sigma Cat. No. A4544) and supplemented with 20 ng/ml recombinant human SCF (Peprotech, Cat. No 300-07) 20 ng/ml recombinant human IL-7 (PeproTech, Cat no 200-07), 10ng/ml recombinant human IL-15 (PeproTech, Cat. No. 200-15), 10 ng/ml recombinant human FMS-related tyrosine kinase 3 ligand (Flt3-Ligand) (PeproTech, Cat. No. 300-19) for the entire duration of NK differentiation, and 5 ng/ml recombinant human IL-3 (PeproTech, Cat. No. 200-03) for the first week of differentiation only. Every 2 to 4 days, a half volume medium change was performed using NK differentiation medium with 2x cytokine concentrations. At the end of the differentiation, on day 28-30, the supernatant containing iNKs was collected into 15 mL conical tubes. A fraction of the cells was kept on ice for flow cytometry as described below to check for purity. The remaining cells were spin down at 300xg for 5 min. at RT and cryopreserved in BiolifeSolutions® CryoStor® CS5 Cell Freezing Medium (StemCell Technologies, Cat. No. 07933) at 5×10^6^ cells per mL per vial according to the manufacturer instructions. For co-culture experiments, iNKs were thawed and cultured overnight in NK differentiation medium without IL-3 at 0.3-0.5x 10^6^ cells per mL in Costar® 6-well Ultra-Low attachment plates (Corning, Cat. No. 3471). Next day, recovered iNKs were collected and counted on a NucleoCounter® NC-202™ Cell Counter (Chemometec) before being used for co-culture experiments.

### Phenotypic characterization of iPSC-derived NK cells via flow cytometry

For every round of iPSC differentiation into NK cells, we first confirmed endothelial-hematopoietic differentiation of embryoid bodies (EBs) cultures via flow cytometry. At least 20 EBs were collected from the initial 96-well differentiation plate. To generate a single-cell suspension from EBs, EBs were enzymatically dissociated with TrypLE™ Express enzyme (Gibco, Thermo Fisher Scientific Cat. No. 12605-010) for an initial 5 minutes incubation at 37°C, followed by shearing of EBs with vigorous pipetting through a P1000 tip, and then a subsequent 2-minute 37°C incubation and second round of vigorous pipetting. Next, the enzymatic dissociation was inactivated by adding 2 mL of Cell Staining Buffer (Biolegend, 420201). The cell suspension was then filtered through a CellTrics™ 50 µm filter (Sysmex, Cat. No. 04-004-2327) and stained with fluorescent antibodies to label cell surface markers. As previously described^74^, EB-derived hemogenic endothelium (HE) was defined as CD34+/CD307high/CD43low/-. FITC anti-human CD34 antibody (Biolegend, 343603), APC anti-human CD43 antibody (Biolegend, 343205) and PE anti-human CD309 (Biolegend, 359903) were used according to the manufacturer instructions. We typically obtained 15-50% HE cells in a 6-day EB culture with the iPSC line used in this study as a source of iNKs. Note that the HE cells yield can vary depending on the iPSC line used. We have typically observed 15-50% HE cells yields across different iPSC lines. For an optimal differentiation into iNKs that renders good yields, EBs should contain at least 30% HE cells. At the end of the differentiation (Day 28-30) iNKs are collected from the supernatant and checked for purity via flow cytometry. iNKs were identified as CD45+CD56+ suspension cells using Pacific Blue™ anti-human CD45 (Biolegend, 368539) and APC anti-human CD56 (Biolegend, 392406). Additionally, cell surface expression of a panel of NK cell markers on CD45+CD56+ iNKs was evaluated via flow cytometry^29,75^ (CD2 (Biolegend, 300207), CD16 (Biolegend, 302007), CD94 (Biolegend, 305506), NKG2A (Miltenyi Biotec, 130-113-566), NKG2D (Biolegend, 320805), NKp30 (Biolegend, 325207), NKp44 (Biolegend, 325107), NKp46 (Biolegend, 331907). For flow cytometry, 0.1 - 0.5 × 10^6^ iNK cells were stained in 50 - 100µL staining buffer containing an antibody cocktail for 20-30 minutes at room temperature in the dark. After staining, samples were washed with 1 mL of Cell Staining Buffer, then spined down for 5 minutes at room temperature. Cell pellets were resuspended in Cell Staining Buffer containing either SYTOX™ Blue (Thermo Fisher Scientific, S34857), with the HE antibody cocktail, or SYTOX™ Green (Thermo Fisher Scientific, S34860), with the NK antibody panel, Dead Cell Stain at 1:1,000 for excluding dead cells in flow cytometry data analysis. Samples were analyzed on an Attune NxT Acoustic Focusing Cytometer (Thermo Fisher Scientific). Flow cytometry data were analyzed with FlowJo™ Software (BD).

### Functional characterization of iPSC-derived NK cells

To determine iNK cells killing capability, we performed a flow cytometry-based assay to quantify iNK cell cytotoxic activity against K-562 cells. K-562 (CCL-243™) or K-562-GFP (CCL-243-GFP™) cells were obtained from The American Type Culture Collection (ATCC®) and cultured according to ATCC’s instructions, in Gibco™RPMI (Thermo Fisher Scientific, 61870036), 10% Gibco™ certified, heat-inactivated FBS (Thermo Fisher Scientific, 10082147), 1x Gibco™ MEM Non-essential amino acids (NEAA) (Thermo Fisher Scientific, Cat. No. 11140050), 1x Gibco™ GlutaMAX™ Supplement (Thermo Fisher Scientific, 35050061), 1x Gibco™ Penicillin/Streptomycin (50 U/mL) (ThermoFisher Scientific, Cat. No. 15070063) 50 µM Gibco™ 2-Mercaptoethanol (Thermo Fisher Scientific, Cat. No. 21985-023), 10mM Gibco™ HEPES (ThermoFisher Cat. No. 15-630-080). To distinguish target tumor cells from iNKs, 10 × 10^6^ K-562 cells K-562 or K-562-GFP were first labeled with 2 μM CellVue™ Maroon (Invitrogen™, 88-0870-16)) according to the manufacturer instructions. For the assay a two-fold dilution series of iNKs starting from 1 × 10^6^ cells per mL to was plated onto Nunc™ 96-Well Polystyrene Conical Bottom MicroWell™ Plate (Thermo Fisher Scientific, Cat. No. 249935) with each dilution plated in triplicate in iNK Assay Medium consisting in a 2:1 (v/v) mixture of DMEM (ThermoFisher Scientific, Cat no 10569010) and Ham’s F-12 (Life Technologies, Cat. No. 31765035), 10% human AB serum (GeminiBio, Cat. No. 100-612), and 25mM Gibco™ HEPES (ThermoFisher Cat. No. 15-630-080). Next, CellVue™ Maroon-labeled K-562 were added onto the assay plate at 0.1 × 10^6^ cells per mL in K-562 culture medium. Wells containing K-562 or iNK alone in iNK Assay Medium were used as controls. The assay plate was then spun down at 120xg for 2 minutes at room temperature and placed in a 37°C incubator for 4 hours. After 4 hours, the plate was spun down at 400xg, for 5 minutes at 4°C and the supernatant was removed. Co-cultures were then washed with 200 μL per well of Annexin V Binding Buffer (Biolegend, 422201) and then stained with 1.6 μg/mL Pacific Blue™ Annexin V (Biolegend) and 20 μg/mL Propidium Iodide (Invitrogen™) in 50 μL Annexin V Binding Buffer per well for 15 minutes at room temperature after which an additional 150 μL per well of Annexin V Binding Buffer was added. Co-cultures and monocultures controls were analyzed on an Attune NxT Acoustic Focusing Cytometer with a CytKick™ Autosampler (Thermo Fisher Scientific). Flow cytometry data were analyzed with FlowJo™ Software (BD). From each well, the percentage of CellVue™ Maroon-labelled cell bodies that was both Annexin V+ and Propidium Iodide+ was quantified as apoptotic K-562.

### Natural killer cells and tumor organoids staining

To set up a co-culture experiment, iNKs were collected from overnight culture after thawing as described above, washed in DPBS (spin 300xg for 5 minutes at room temperature) and counted prior to labeling. iNKs were then labeled with 20 μM eBioscience™ Cell Proliferation Dye eFluor™ 450 (Invitrogen™) at 2000 iNKs/μl DPBS for 10 minutes at 37°C. The staining reaction was quenched with 5x the volume of the staining suspension using chilled RPMI (Life Technologies, Cat. No. 61870036) containing 10% heat-Inactivated fetal bovine serum (HI-FBS; Gibco Cat. No. 10082147) (RPMI/10% HI-FBS) for five minutes on ice. Stained iNKs were then spun down at 450xg for 5 min. at room temperature and washed three times with RPMI/10% HI-FBS. The labeled iNK cell pellet was then resuspended in iNK co-culture media, consisting of the iNK differentiation media and containing 2x concentration of growth factors and cytokines as follow: 40 ng/ml recombinant human SCF (Peprotech, Cat. No 300-07), 40 ng/ml recombinant human IL-7 (PeproTech, Cat no 200-07), 20ng/ml recombinant human IL-15 (PeproTech, Cat. No. 200-15), and 20 ng/ml recombinant human Flt3-Ligand (PeproTech, Cat. No. 300-19), counted, and kept on ice until plating.

For PDOSs staining, organoids were expanded for 6 days in standard culture conditions and extracted intact as fully formed organoids. To extract the organoids from the Cultrex® BME domes, the culture media was aspirated and replaced with cold Corning® Cell Recovery Solution (Fisher Scientific, Cat. No. CB-40253) containing 10 µg/mL DNase I (Sigma-Aldrich, Cat. No. DN25), and 10 µm Y-27632 ROCK inhibitor (AbMole, cat. No. M1817). We then incubated the plate at 4°C for 30 min. in a rocking platform. Organoids were then collected and washed three times with 2-3x the collected volume using cold Gibco™ Advanced DMEM/F12 (ThermoFisher Scientific, Cat. No. 12634-028) containing 0.1% BSA (Gibco, Cat. No. 15260-037), 20 µg/mL DNase I (Sigma-Aldrich, Cat. No. DN25) and 10 µm Y-27632 ROCK inhibitor (AbMole, cat. No. M1817). Organoids resuspended in the washing media were placed at 37 °C for 10 min. then spun down at 200-250xg, 4-8°C for 5 min. After washes, the organoid pellet was resuspended into the same media without BSA for counting for which we used Keyence hybrid cell count analysis software (Supplementary Fig. 2). Organoids were then labeled with 15 μM CellTracker™ Orange CMRA (ThermoFisher) at a concentration of 50×10^3^ organoids/mL in an Eppendorf® Protein LoBind tube (Millipore Sigma, Cat. No. EP0030122356) at 37°C for 45 minutes, with gentle mixing every 15 minutes. At the end of 45-minute incubation, organoids were spun down at 200xg for 1 min at room temperature and washed three times with 1.5x the staining volume using tOVA media. Labeled organoids were resuspended in tOVA media, filtered through a 70 µm Fisherbrand™ Sterile Cell Strainer (Fisher scientific, Cat. No. 22-363-548), counted as described above and kept on ice until plating.

### Patient-derived ovarian cancer organoids and iNKs co-culture

For co-culture experiments, we used PhenoPlate™ 96-well microplates (Revvity Health Sciences, Cat. No. 6055300) with freshly coated wells using 100 µL of 1:1 (v/v) DMEM (ThermoFisher Scientific, Cat no 10569010):Ham’s F 12 (Life Technologies, Cat. No. 31765035) containing 10% Cultrex® BME for at least 2 hours at 37ºC before plating. The coating media was aspirated from the assay plates right before plating. In this study, we used 20:1 Effector (iNKs) to Target (organoid units) (E:T) ratio *per* well. To standardize the plating and minimize variability across experiments, the labeled iNKs and PDOs suspensions were brought up to 300K/ml and 15K/ml respectively using the same media they were resuspended on. For PDOs we also added Cultrex® BME at 10% final concentration, DNase I (Sigma-Aldrich, Cat. No. DN25) at 20 µg/mL final concentration, and CellEvent™ Caspase-3/7 detection reagent (ThermoFisher Scientific, Cat. No. C10423) at 10 µM final concentration. iNKs and PDOs solutions were then mixed at 1:1 (v/v) ratio, placed on ice and gently pipetted up and down twice prior to plating. 200 µL of the iNK/PDO solution was added to each well resulting in 30,000 iNKs /1,500 PDOs per well. For monoculture controls, the PDOs solution was mixed with iNK co-culture media containing 2x concentration of growth factors and cytokines as described above. The assay plates were then spun down at 50 g for 1 minute at room temperature and placed in Opera Phenix™ high-content imaging screening system (Revvity) for imaging.

### Live-cell imaging of PDOs and iNKs co-cultures

Plates were imaged using Opera Phenix™ high-content imaging screening system (Revvity) at 37ºC and 5% CO2. Images were acquired every 15 minutes for up to 11 hours. Assay plates were stored in an automated incubator at 37ºC and 5% CO2 (Cytomat, Thermofisher) between each acquisition time point. Images were acquired in confocal mode with a 10x (Air, NA 0.3) or 20x objective (Water, NA 1.0) using the following excitation wavelengths: 388 nm to acquire eFluor-iNKs, 488 nm to acquire Caspase-3/7 reagent, and 561 nm to acquire CMRA-PDOs respectively. The entire well well was imaged using (21 fields in total) with Z-stack images acquired within a range of 103.6 µM, with 7.4 µM between each plane for a total of 15 planes. Analyses were performed using the Opera Phenix™ Harmony software, version 4.9.

### Image analysis

Image analysis was performed using the Harmony High-Content Imaging and Analysis Software. Z stack images were overlayed to create a max projection. Organoids were defined as objects with a surface area over 314 µm^2^ (corresponding to > 20 µm in diameter) exhibiting CMRA fluorescence at a minimum of 1000 intensity unit (CMRA positive). Caspase-3/7 activation was defined as a minimum signal of 2500 intensity unit. To quantify organoid treatment response, the percentage of the total organoid area (i.e. CMRA positive) for each individual organoid overlapping with positive Caspase-3/7 signal was quantified at each time point.

### Bayesian modelling of organoid apoptosis

To investigate resistant and sensitive organoid populations, we used brms 2.22.0 to fit a Bayesian two-component Beta mixture model, where the apoptosis outcome is distributed as a mixture of two Beta distributions and the mixture proportion *θ*_2_ is predicted based on the patient covariate *p*_*i*_.

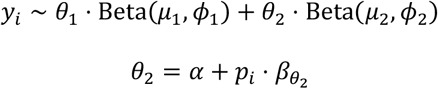

The Beta distribution was parametrized using a mean (*µ*) and a precision (*ϕ*) parameter as implemented in brms. The softmax transformation was applied to the linear predictor terms for the mixture proportions to ensure that *θ*_1_ and *θ*_2_ were valid probabilities summing to 1. The priors on the mixture model parameters were:

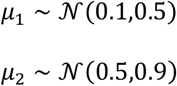

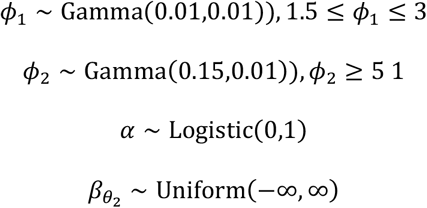

The softmax transformation was applied on the linear predictor terms for the mixture weights to ensure that *θ*_1_ and *θ*_2_ are valid probabilities in the interval (0,1) that sum to 1. Before model fitting, to avoid apoptotic surface area values of 0 and 1, values were normalized to the exclusive 0-1 range^76^. The model was fit using 4 chains with 4,000 iterations per chain and convergence was determined based on R-hat values <1.01^77^ and effective sample size (ESS) >=1000^78^. Point estimates and credible intervals (CI) for parameters were calculated based on the fitted model. We used posterior predictions from the fitted model to calculate the difference in resistant organoids between the two patients using the compare_levels function from the tidybayes package^79^. We calculated posterior probabilities of mixture component memberships across observations with brms and determined a threshold for membership in the resistant organoid group. This threshold was then applied across experimental timepoints to compare mean apoptosis between non-apoptotic and susceptible apoptotic organoids.

To fit the percentage of apoptotic organoids over time based on this threshold and identify the asymptote representing the persistent percentage of resistant organoids, we compared logistic, Hill and Gompertz models. These non-linear models were fit to the data with brms as described above, using a Beta family error model. The logistic model was assigned broad priors for the midpoint *t*_0_ and the slope *k*:

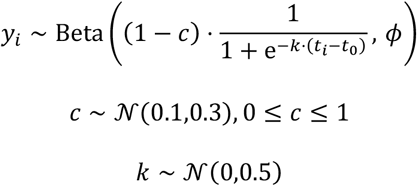

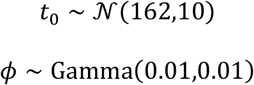

The Hill model was assigned similar priors for the midpoint *K*_*m*_ and the slope *α*:

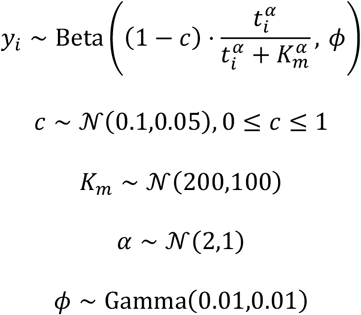

Finally, for the Gompertz model, similar priors were assigned for the displacement along the x-axis *b* and growth rate *d*:

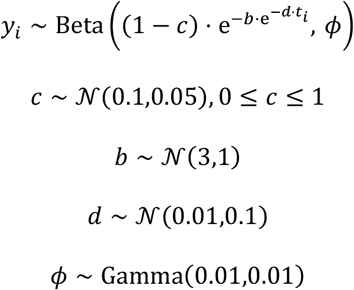

All models were assigned the same prior for the upper asymptote *c* and all models had the same prior for the precision of the Beta error model *ϕ*. Fitted models were compared using loo and the report 0.6.1 package in R and a difference >4 in expected log predictive density (ELPD) was used as a threshold for notable differences between models^80^.

To determine the two distinct distributions of small and large organoids at the experimental endpoint, a Gaussian mixture model was used:

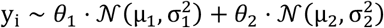

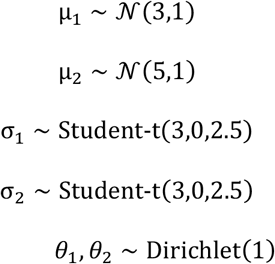

Fitting parameters for brms were set as above. The mixing proportions *θ*_*i*_ and *θ*_1_ were normalized to sum to 1. The posterior probabilities of memberships in the small and large organoid group were computed using the pp_mixture function in brms. Observations were then assigned to the group with the larger posterior probability of membership.

